# Rescuing error control in crosslinking mass spectrometry

**DOI:** 10.1101/2023.11.28.568978

**Authors:** Lutz Fischer, Juri Rappsilber

## Abstract

Crosslinking mass spectrometry is a powerful tool for studying protein-protein interactions under native or near native conditions in complex mixtures. By help of novel search controls, we show that measures that aim to improve the number of identifications based on heuristic considerations can undermine error estimation in non-obvious ways. The relationship between decoys and false positives is very sensitive to the injection of information that favour likely correct matches. We identify a wider challenge in crosslinking data analysis tools with maintaining the decoy-false positive relationship, and exemplify this with the filtering of results based on the information of which proteins can be observed as having reacted with the crosslinker. Without correcting for this problem, we could identify an average of 260 interspecies protein-protein interactions in 16 analyses, “solidly” suggesting groundbreaking biological connections that don’t actually exist. We also show how data analysis procedures can be tested and modified to rescue the decoy-false positive relationship. The importance of this relationship for reliably identifying protein-protein interactions cannot be overstated.

## Introduction

Crosslinking mass spectrometry (MS) has emerged as a powerful approach for studying protein-protein interactions in native or near-native conditions^1,2^. This technique involves the addition of a crosslinker to a protein sample to covalently capture protein interactions, followed by the digestion of the sample and the identification of the resulting crosslinked peptide pairs using mass spectrometry. The identification of crosslinked peptides is confounded by many factors, which notably includes a large database search space. In addition, crosslinked peptide pairs originating between distinct protein sequences (protein heteromeric links^3^) tend to be less abundant than crosslinks between peptides within one protein sequence (self-crosslinks) and suffer from higher random matching (due to their lower abundance in the sample and more theoretically possible combinations in the search space), leading to lower scores and higher noise levels. Consequently, distinguishing true protein heteromeric crosslinks from false positives can be difficult, limiting the sensitivity of crosslinking MS identifications.

It is tempting to turn to information sources beyond the individual crosslink-spectrum matches (CSMs) to improve the number of identified protein heteromeric crosslinks. One such method, called mi-filter^4^ (Figure 1), aims to reduce the false positives among the protein heteromeric matches by only considering matches between proteins that are also observed with self-crosslink matches or with linear crosslinker modified peptides (monolinks). The underlying assumption is: any protein seen in a protein heteromeric crosslink should also be observed with modified linear peptides or self-crosslinks. This is based on the general observation that modified linear peptides, and also self-crosslinks, tend to be more readily observed due to their higher abundance than protein heteromeric crosslinks. Leveraging these observations correctly, though, might be more difficult than initially meets the eye, and doing so wrongly would destroy the validity of the error models underlying any decoy-based false discovery rate (FDR) approach.

**Figure 1:**
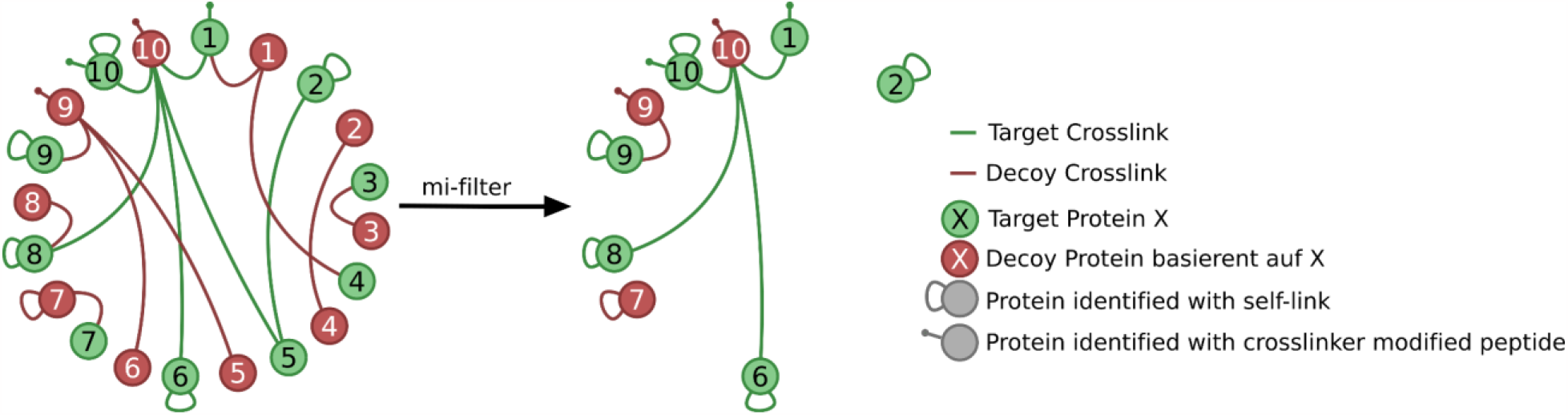
When using the mi-filter, only proteins that are observed with crosslinker modified linear peptides or as part of a self-crosslink are kept and considered as observable in heteromeric protein-protein interactions.

Typically, in crosslinking mass spectrometry, decoy matches are taken as known false positives and used to estimate the amount of unknown false positives among the target matches. This allows results to be filtered to a defined level of confidence^5–10^. This is expressed in the form of a false discovery rate (FDR). Decoy matches refer to matches to a protein or peptide that was searched but is known to be absent from the sample. This is usually accomplished by searching either reversed target proteins^5^ or randomly generated proteins^11^, where each decoy protein represents an equivalent target protein. Importantly, the amount and score distribution of decoy matches provide a model for the amount and distribution of false positive target matches. By comparing the number of decoy matches to the number of target matches the FDR can be estimated and in turn a score threshold can be defined to filter the search results to a specified FDR.

Here, we demonstrated that mi-filter and similar approaches disadvantage decoys over targets and thereby break the ability of decoys to model the noise among the target hits. Hence, we can’t use the remaining decoys to assess the confidence of the results. Also, we show how to correctly apply this kind of filter while keeping the decoy-false positive equivalency.

## Methods

The datasets used here were downloaded from ProteomeXchange^12^. The *Mycoplasma pneumoniae* dataset was taken from Pride ID PXD017711^13^, specifically the raw files for the strong cation chromatography (SCX) fractions 11 to 14. The *E. coli* dataset was taken from JPOST ID JPST000845^10^, here the DSSO raw files for SCX fraction 18, 20, 22 and 24. The 26S proteasome dataset comprises the trypsin-only raw files from Pride ID PXD008550^14^.

The raw files were converted to mgf-files with ProteoWizard (version 3.0) msConvert^15^. Search parameters used were: crosslinker DSSO for *M. pneumoniae* and *E. coli* dataset and BS3 for the 26S proteasome dataset with specificity for Lysine, Serine, Threonine, Tyrosine, and protein n-terminal with a bias towards Lysine and n-terminal, fixed modification of carbamidomethylation of Cysteine, variable modification of oxidation on Methionine and hydrolyzed and amidated crosslinker modifications on K, S, T, Y, and protein N-termini as linear only modifications. Non-covalent interactions were considered as part of the search as well but ignored during data analysis.

The *E. coli* and *M. pneumoniae* datasets were searched against a database of all *E. coli* proteins (UniProt proteome UP000000625 from 19/01/2023) and all *M. pneumoniae* proteins (UniProt proteome UP000000808 from 16/01/2023). For data analysis, each *E. coli* SCX fraction was paired with each *M. pneumoniae* SCX fraction and the average of all combinations plotted. Searching against a joined database provides a decoy-independent set of known false positives, and hence a control if decoys correctly reflect false matches among the targets. Any match has to be wrong that is between *E. coli* and *M. pneumoniae* proteins but also any match of an *M. pneumoniae* protein to an *E. coli* spectrum and inversely any match of an *E. coli* protein to an *M. pneumoniae* spectrum.

The 26S proteasome dataset was first searched with MaxQuant^16^ version 1.6.17 against complete *Saccharomyces cerevisiae* proteome (UniProt proteome UP000002311 from 06/02/2023) resulting in a list of 1073 non-contaminant target protein groups. Of each, the first protein was taken as a representative. This list was sorted by iBAQ value and split in three sets. The non-identified proteins from the original FASTA file were split into 10 subsets. For the crosslinking MS data analysis, xiSEARCH was run first against increasing numbers of present proteins in order of abundance. To test the effect of the filter with non-present proteins, the search was repeated against all of these identified proteins + increasing sets of non-identified proteins. For data analysis, each fraction was individually searched and filtered.

To enable a valid FDR calculation, the datasets were filtered to only accept the highest scoring match for any given combination of peptide pairs, precursor charge state modifications and linkage-site.

## Results

### Classes of false positives

In the context of protein-based filters in crosslinking MS, we can describe two different types of false positive matches within the realm of target matches:

#### False Positive Group 1

This category encompasses random matches that involve at least one protein that is not observable as part of a genuine crosslink. This situation arises either because the protein is absent from our sample or due to practical factors like low protein abundance, rendering it undetectable as part of a crosslink. If such a protein is nevertheless identified by being matched to a spectrum, it constitutes a false positive. Of course, one does not know which specific matches this applies to. However, one knows that this error occurs.

#### False Positive Group 2

This category encompasses random matches between protein pairs where both proteins are observable as part of genuine crosslinks. This means that these proteins are detectable as interacting with each other or with one or more other proteins. Being part of a genuine crosslink does not mean that a protein can not also be matched in a false positive peptide-spectrum match. In other words, a random match may still occur involving a protein with multiple correctly identified crosslinks. For example, if two protein pairs, AB and XY, truly existed independently of each other in a sample, one could still identify false matches between them, i.e., AX, AY, BX, BY, AA, BB, XX, and YY.

In the absence of any rescoring or filtering, decoys model both of these false positive groups. However, this conventional approach becomes inadequate when a filter is introduced to distinguish between these two distinct groups, as it necessitates a more nuanced modelling strategy.

### Theoretical weakness of the mi-filter algorithm

The mi-filter makes a heuristic decision to limit crosslink-observable proteins to only those proteins that were observed with self-crosslinks or linear modified peptides. Other heuristics have been employed by similar approaches. These include restricting the search to those proteins that are in the sample, i.e. can be identified by any peptide^5,17^ or those proteins that are identified with a certain abundance^10,14^. Asking for self-crosslinks or linear modified peptides relates to the abundance criterion, as these peptides tend to have lower abundance than linear unmodified peptides but higher abundance than protein heteromeric crosslinks. One also adds the observation that peptides of those proteins actually reacted with the crosslinker, although, it is unclear if this is important. In this way, the mi-filter excludes proteins that are likely not observable as being crosslinked with other proteins. This reduces noise matches, predominantly reducing false positives of group 1. Note that the mi-filter also excludes small proteins that are difficult to detect, making them even more difficult to detect^10^.

However, the mi-filter, being applied as a filter after the search and not before, requires some considerations of how target and decoy proteins are affected. The decoys that pass the mi-filter now model proteins that pass the filter falsely, i.e., are not observable as self-crosslink or crosslinker modified. In line with the initial hypothesis, these can be assumed to be not observable as part of a heteromeric link either. As a result, after filtering the protein heteromeric matches, any decoys can now only represent false positive matches from the false positive group 1. However, matches from false positive group 2 remain present. Consequently, any FDR calculation relying on decoys will underestimate the total error.

### The mi-filter leads to severely underestimating the error

To assess if the mi-filter leads to detectable underestimation of error, we constructed a test case of two sets of proteins that are crosslinked only within each set but permitting “identifications” of crosslinks also between the sets of proteins. This allowed us to reveal a large proportion of false protein pairings of the type AX, AY, BX, and BY in the example above. Entrapment databases are often used for testing the accuracy of an error assessment approach^10^. Unfortunately, a simple entrapment search only provides ground truth for the absence of proteins, and in effect can only test if false positive group 1 (crosslinks involving non-present proteins) is modelled. For this reason, we took instead the data of two separate large-scale crosslink investigations, from *E. coli*^*10*^ and *M. pneumoniae*^*13*^, and searched against a combined database of *E. coli* and *M. pneumoniae* proteins. In this way, the *M. pneumoniae* proteins become the entrapment database for the *E. coli* data and the *E. coli* proteins the entrapment database for the *M. pneumoniae*. Importantly, both species contain observable proteins that at the same time will be visibly false positive when they are matched to spectra of the other species or in a pair together with a protein of the other species.

Looking at all matches passing a minimal score cut-off (xiSEARCH score >= 5), the decoy-based estimate of false positives exceeds the number of observed impossible matches (Fig. 2A). This is expected, since the possible matches (i.e., *E. coli* protein pairs matched to *E. coli* spectra and *M. pneumoniae* protein pairs matched to *M. pneumoniae* spectra) will of course also contain wrong results (false positives). After applying the mi-filter, few decoys remain, resulting in a significantly reduced number of estimated false positives. However, the number of impossible matches, crossing the dataset boundaries, is notably higher than they should if the FDR estimate were correct. There are >5 times more impossible matches than estimated false positive matches based on decoys. Thus, the actual FDR of the mi-filtered results has to be larger than 33%, while the decoys state an apparent FDR of 6.2% at the crosslink-spectrum match level. This means that a large part of the alleged gain from the mi-filter is only the result of hiding the actual error (Figure 2A red hatched area). If one were to now apply an FDR-based cut-off such as 1% or 2% decoys, this would leave 8.6% or 15.9% impossible matches respectively and thus a substantially higher error in the reported data than expected. In consequence, we could mistakenly “discover” extensive protein-protein interactions between *E. coli* and *M. pneumoniae* (in our 16 test analyses, on average 260 interspecies PPI based on 2% CSM-FDR or 67 at 2% PPI-FDR), suggesting groundbreaking biological connections that don’t actually exist. These findings would stem from a serious data analysis mistake.

**Figure 2:**
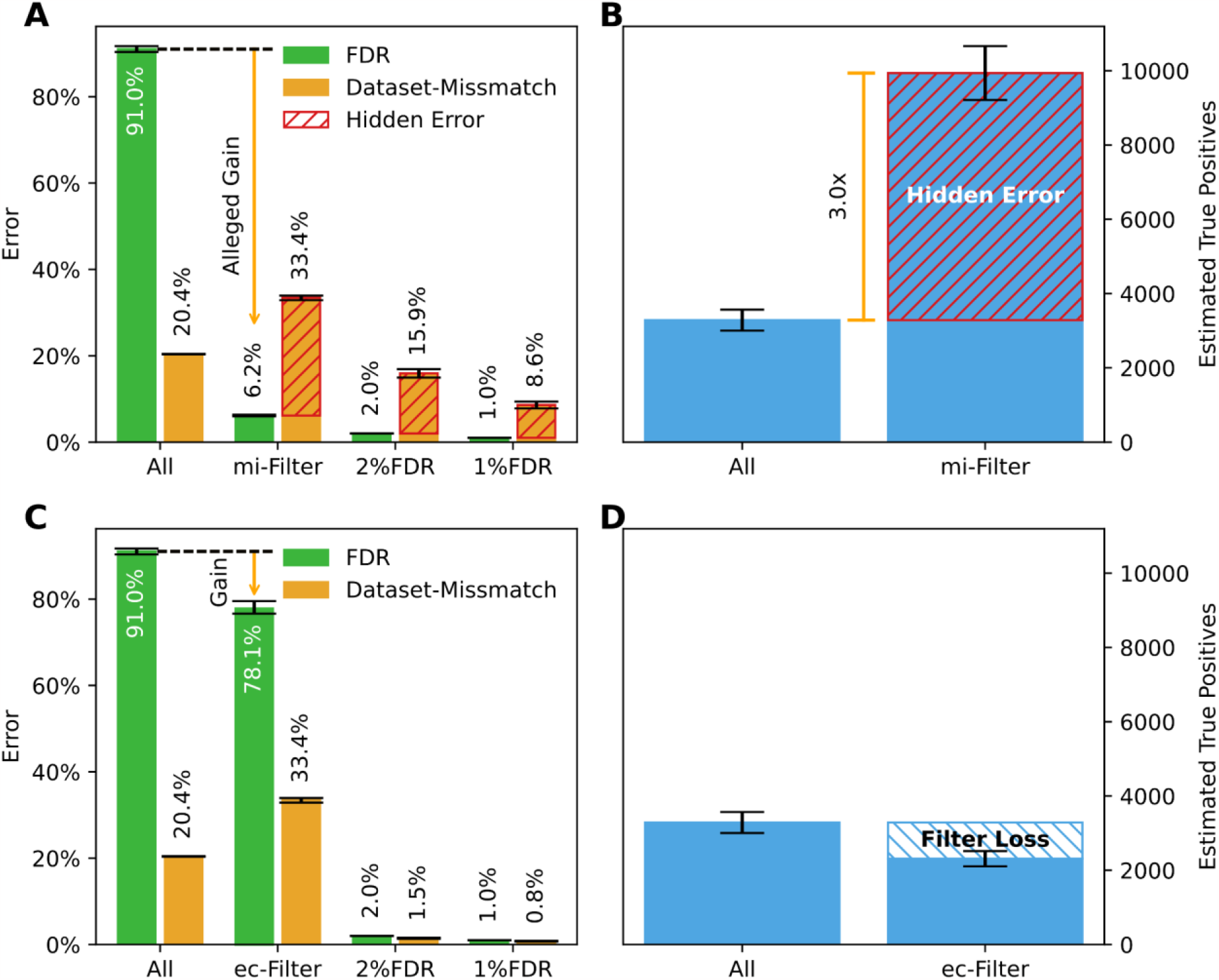
Effects of applying mi-filter and corrected ec-filter. A) decoy based FDR estimate (green) and fraction of dataset mismatches (matches to *E. coli* spectra that involve *M. pneumoniae* proteins and matches to *M. pneumoniae* spectra that involve *E. coli* proteins) with a minimum score cut-off of 5 (All), after applying the mi-filter or after mi-filter and a 2% and 1% FDR cut-off. The mi-filter results in a 15.9% dataset mismatch rate at a 2% FDR cut-off. The shaded areas indicate the known false positives not modelled by decoys. B) Decoy based estimate of assumed true positives among results for either just applying a minimum score cut-off (left) or after applying the score cut-off and mi-filter (right). The shaded area indicates the number of impossible true positives. C) Same as A, but with the corrected ec-filter. D) same as B) but with the corrected ec-filter The shaded area indicates how many true positives are filtered out by employing the ec-filter.

### A universal test for error estimates being affected by filters

A more universal approach to assess if a filter breaks the estimation of false positives would be to assess how the approach affects the estimated true positives, instead of false positives. The number of true positives in the sample can be estimated as

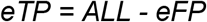

With eTP being the number of estimated true positives, eFP being the estimated false positive matches and ALL being the total number of matches in which the amount of false and true positives are to be estimated.

As we here use decoys to model false positive crosslink matches, *eFP* turns into^6,8^

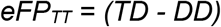

*With* eFP_TT_ representing the estimated number of false positives among the target-target matches, TD the number of matches involving one target and one decoy part, and DD the number of matches that involve only decoys. The possibly surprising subtraction of DD from TD in this formula is the result of search space considerations^8^.

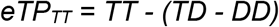

With TT being the number of matches that match exclusively the target database.

We assume that the way to estimate the number of false positives (here decoy based), and in extension the estimate of all true positives, is in fact correct. Then this gives an upper bound of how many true positives can at most be identified. Hence, the total number of similarly estimated true positives *after* applying any filter should never exceed that present in the unfiltered dataset. In actuality, almost all correct filters will also reduce the total number of true positives in a dataset. However, for the mi-filter, the number of true positive crosslink-spectrum matches (CSMs) estimated after applying the mi-filter on CSMs exceeds that present in the unfiltered dataset by a factor of 3.0 (Fig. 2B). This indicates a serious overestimation of true positives and consequently an underestimation of false positives. Note that this test can only reveal that something is wrong. Passing this test is no proof of correctness. Also, this test can only work if all data used directly as input are compared to all data directly passing the filter in question. As soon as additional steps are applied (e.g. FDR, score cut-offs, quality metrics…) the result of this test will be obscured.

### How to restore the decoy - false positive relationship

Proteins that are observable as part of a protein heteromeric crosslink are likely also observable via self-crosslinks^10^ or linear crosslinker-modified peptides^18,19^. Hence, filtering protein heteromeric matches to proteins that are seen as part of a self-crosslink or as linear crosslinker modified peptide should reduce noise and hence might improve the detection of protein heteromeric crosslinks.

As we show, however, the implementation of these ideas by the mi-filter disrupts the decoy-to-false-positives relationship by selectively retaining a specific subset of decoy proteins that represent only non-present and non-crosslinkable proteins. To accurately estimate the number of false positives after applying such a filter, it is necessary to include an additional set of “acceptable” proteins. We maintain the relationship between decoy and false matches by considering the decoy complement for each protein identified with self-crosslinks or linear crosslinker modified peptides (Fig. 3). That means, for every target protein that successfully passes the initial filter, any crosslink involving the corresponding decoy protein derived from that target protein is also accepted. For instance, if protein A is identified, we will accept the reversed form of protein A as passing the filter as well. Likewise, for decoy proteins, we accept the original target protein. By doing so, we maintain a balanced target-decoy relationship for heteromeric proteins even after the filter. With this modification, both tests in Figure 2C/D show no contradiction.

**Figure 3:**
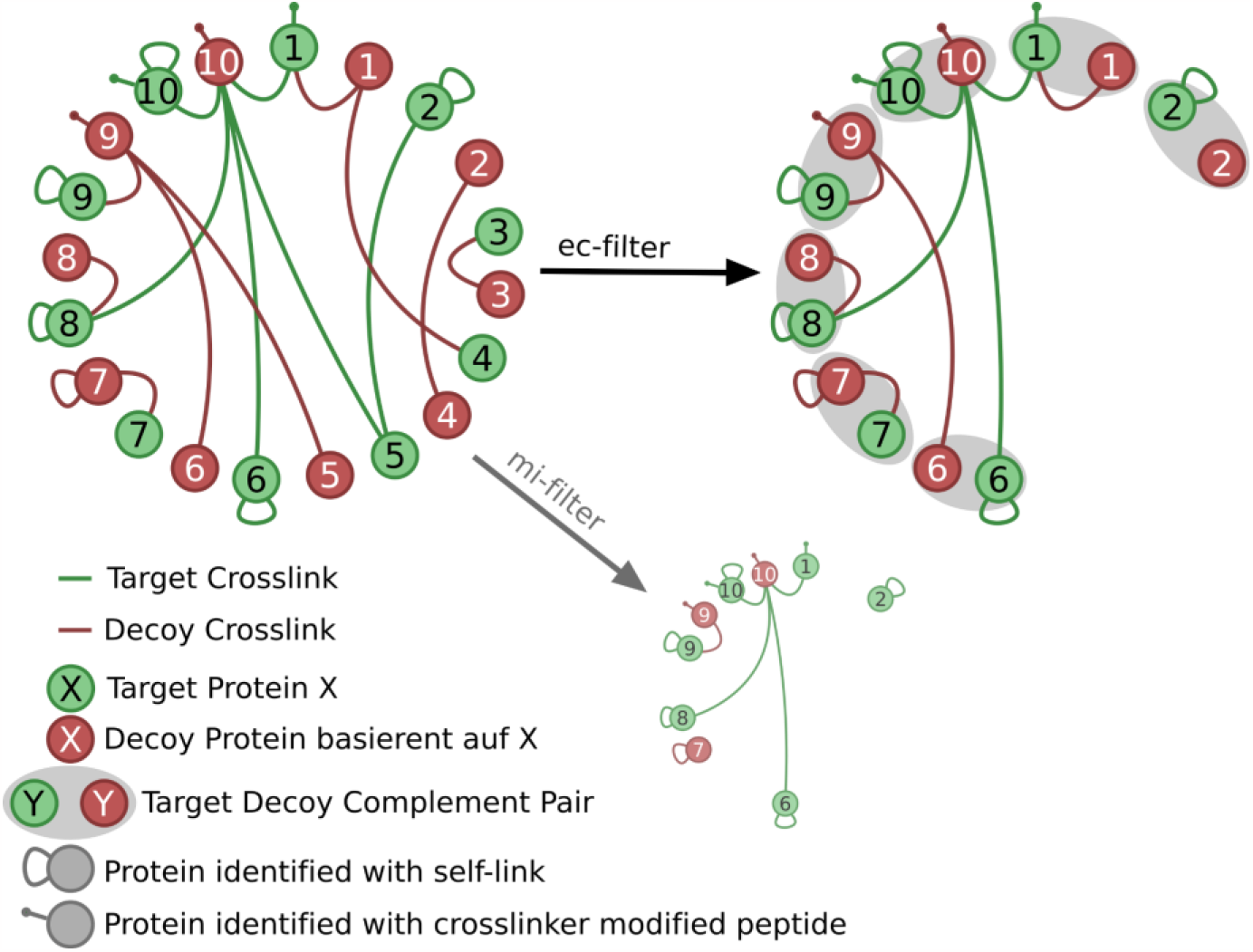
ec-filter filters by presence of self-crosslinks or crosslinker modified peptides but treats target and decoy proteins as a pair, and if one is passing the complement is always passing as well. Small circle is the mi-filter representation for comparison.

Having established a filter (ec-filter - expected crosslinked proteins filter), that maintains decoys as a model of both false positive groups, we then evaluated the extent to which the filter improves the number of protein heteromeric matches. For this, we searched a BS3 crosslinked 26S proteasome dataset with increasing numbers of proteins. First, we used three increasingly larger databases comprising only proteins identified as part of a standard MaxQuant search and then searched against databases additionally supplemented with proteins not previously identified.

The ec-filter on its own (solid orange line vs. solid blue line) can be helpful, especially in those cases where many proteins are searched that are unlikely to be observed as part of a crosslink (Fig. 4). Note that xiFDR offers an option to boost results by using a different set of prefilter (filtering on lower-level FDRs and additional subscores)^8^. When this is employed, the gains of the ec-filter dwindle. The difference between using the ec-filter + boosting and just using boosting is relatively small, with only a slight advantage observed for the ec-filter in case of a database with large excess of proteins that are not actually present in the sample.

**Figure 4:**
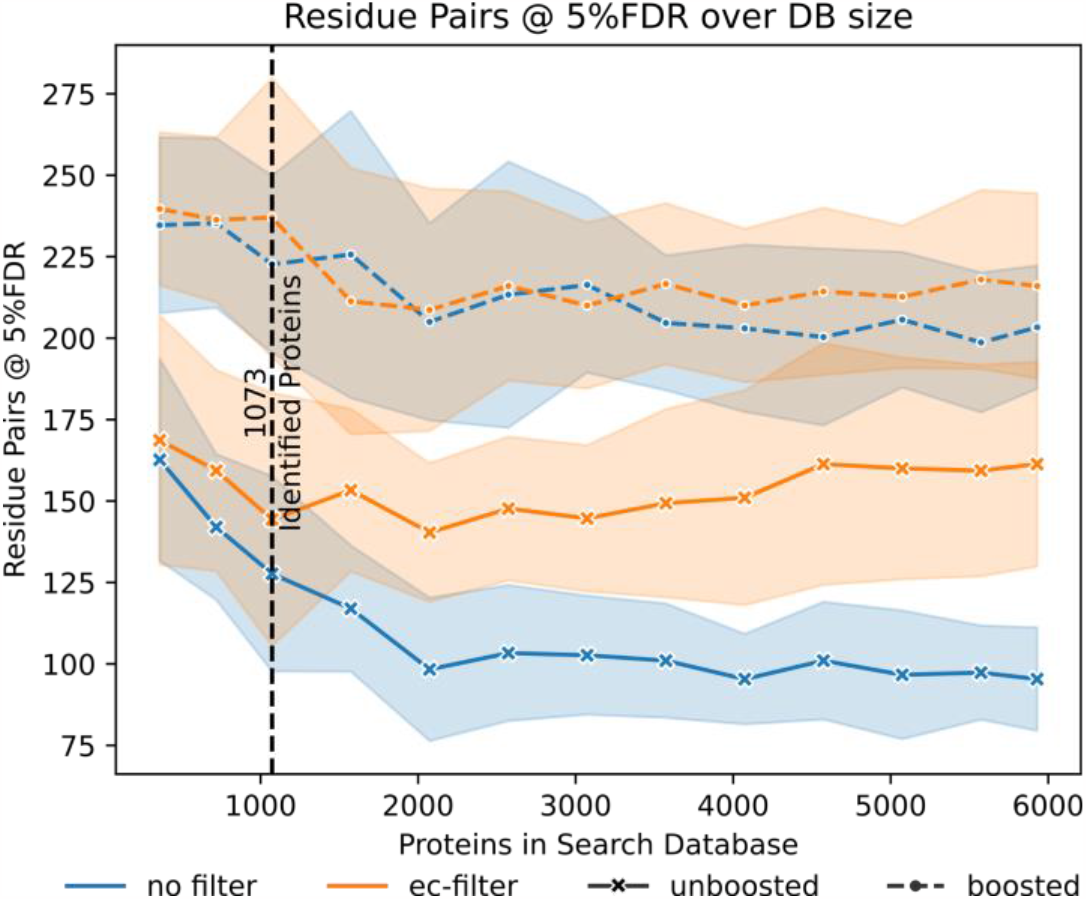
Results of the xiFDR 2.2 implementation of the ec-filter at a fixed 5%-residue-pair FDR, depending on initially searched database size. Data plotted are the average results among three fractions searched individually against the same database. The coloured area represents the standard error.

## Discussion

Our study highlights the need to be aware of how filters affect the decoy-false positive relationship and to take steps to correct for any changes in this relationship. The recently published mi-filter destroys the balanced decoy-target relationship. With the ec-filter we show a filter based on the same observations of linked propensities for proteins to be detected as part of a protein heteromeric crosslink when having been identified with self-crosslinks or modified linear peptides. This ec-filter can improve the number of protein heteromeric matches at a given confidence without underestimating the number of false positives.

The relevance of our findings goes beyond the realm of post-search filtering. Generally, using information about individual proteins for the assessment of protein pairs, both derived from within the search and derived from external sources, has to be done with great care. Search engines or post-processing tools such as ECL-PF^20^ and CRIMP 2.0^21^ utilise protein self-crosslinks to assign scores or confidence values to protein heteromeric matches. XLinkProphet^22^ uses information about individual proteins being present to rescore matches. It is not always clear from the available descriptions what is done exactly. The developers now have the tools, however, to ensure correct handling of information: When a protein is assigned a higher confidence level, the same treatment must be applied to its corresponding decoy complement, following the principles of the ec-filter.

We would like to emphasise the importance of exercising caution when incorporating external information into data analysis processes. This holds true also within the data analysis process. For example, if both residue-pair-level false discovery rate (residue pair FDR) and protein-protein interaction-level false discovery rate (PPI-FDR) are used, it is crucial to complete the assessment of residue-pair FDR before proceeding with PPI-FDR (assuming the PPI-level estimate is based on the residue-pair-level estimate). More generally, if error assessment is done consecutively on several levels of consolidation, where the output of an FDR filter is the input of another FDR filter, then it has to be done in the order of the level of consolidation. I.e. CSM before peptide pairs before residue pairs before PPIs. Engaging in the reverse order, i.e., conducting PPI-FDR first and subsequently addressing residue pair FDR with the remaining decoys, can inadvertently lead to the same pitfall as observed in the mi-filter approach—namely, considering only a subset of the remaining errors, which can compromise the overall accuracy of the analysis.

We have implemented all our current understanding of correct decoy-based error handling in crosslinking, including the ec-filter, in the open source, error-estimation software xiFDR (https://www.rappsilberlab.org/software/xifdr/) version 2.2 (Supporting Figure S1). We would like to encourage any changes or additions to FDR estimation to be communicated openly to its GitHub repository (https://github.com/Rappsilber-Laboratory/xiFDR). Findability, accessibility, interoperability, and reusability (FAIR) relate only to data access. Data use critically depends on trust, and trust depends on robust error handling. This can only be achieved as a joint effort of the crosslinking mass spectrometry community.

## Acknowledgements

We would like to gratefully acknowledge Dr. Colin Combe and Dr. Andrea Graziadei for fruitful discussions regarding the manuscript and FDR in crosslinking.

---

**Supporting Figure S1:**
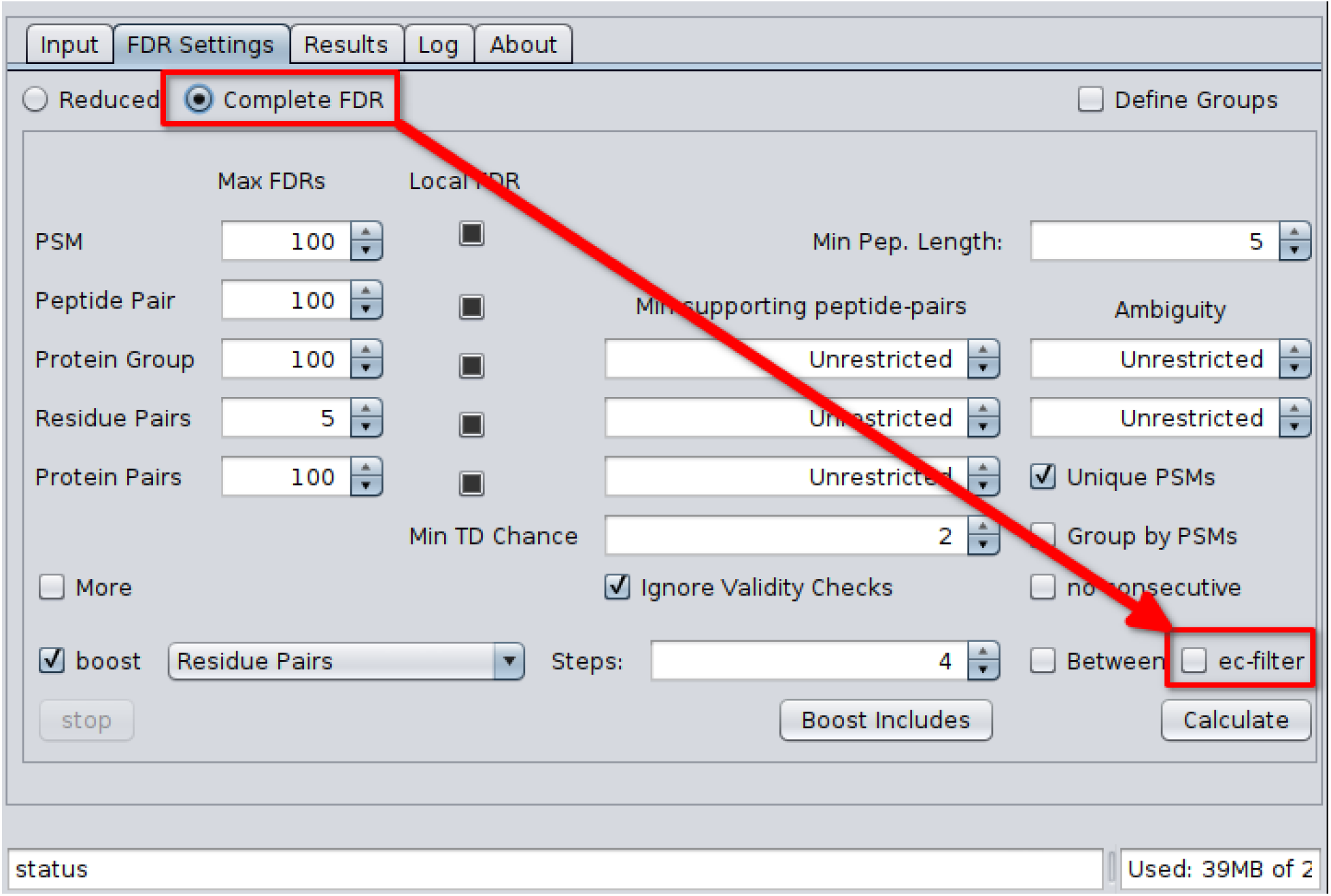
To use the ec-filter the complete settings need to be used and the ec-filter checkbox ticked.

